# Two novel loci underlie natural differences in *Caenorhabditis elegans* macrocyclic lactone responses

**DOI:** 10.1101/2021.01.14.426644

**Authors:** Kathryn S. Evans, Janneke Wit, Lewis Stevens, Steffen R. Hahnel, Briana Rodriguez, Grace Park, Mostafa Zamanian, Shannon C. Brady, Ellen Chao, Katherine Introcaso, Robyn E. Tanny, Erik C. Andersen

**Affiliations:** Molecular Biosciences, Northwestern University, Evanston, IL; Interdisciplinary Biological Sciences Program, Northwestern University, Evanston, IL

## Abstract

Parasitic nematodes cause a massive worldwide burden on human health along with a loss of livestock and agriculture productivity. Anthelmintics have been widely successful in treating parasitic nematodes. However, resistance is increasing, and little is known about the molecular and genetic causes of resistance. The free-living roundworm *Caenorhabditis elegans* provides a tractable model to identify genes that underlie resistance. Unlike parasitic nematodes, *C. elegans* is easy to maintain in the laboratory, has a complete and well annotated genome, and has many genetic tools. Using a combination of wild isolates and a panel of recombinant inbred lines constructed from crosses of two genetically and phenotypically divergent strains, we identified three genomic regions on chromosome V that underlie natural differences in response to the macrocyclic lactone (ML) abamectin. One locus was identified previously and encodes an alpha subunit of a glutamate-gated chloride channel (*glc-1*). Here, we validate and narrow two novel loci using near-isogenic lines. Additionally, we generate a list of prioritized candidate genes identified in *C. elegans* and in the parasite *Haemonchus contortus* by comparison of ML resistance loci. These genes could represent previously unidentified resistance genes shared across nematode species and should be evaluated in the future. Our work highlights the advantages of using *C. elegans* as a model to better understand ML resistance in parasitic nematodes.

**Author Summary:** Parasitic nematodes infect plants, animals, and humans, causing major health and economic burdens worldwide. Parasitic nematode infections are generally treated efficiently with a class of drugs named anthelmintics. However, resistance to many of these anthelmintic drugs, including macrocyclic lactones (MLs), is rampant and increasing. Therefore, it is essential that we understand how these drugs act against parasitic nematodes and, conversely, how nematodes gain resistance in order to better treat these infections in the future. Here, we used the non-parasitic nematode *Caenorhabditis elegans* as a model organism to study ML resistance. We leveraged natural genetic variation between strains of *C. elegans* with differential responses to abamectin to identify three genomic regions on chromosome V, each containing one or more genes that contribute to ML resistance. Two of these loci have not been previously discovered and likely represent novel resistance mechanisms. We also compared the genes in these two novel loci to the genes found within genomic regions linked to ML resistance in the parasite *Haemonchus contortus* and found several cases of overlap between the two species. Overall, this study highlights the advantages of using *C. elegans* to understand anthelmintic resistance in parasitic nematodes.

## Introduction

Parasitic nematodes pose a significant health and economic threat, especially in the developing world [1–3]. These infections increase morbidity and exacerbate the deleterious effects of malaria, HIV, and tuberculosis [4]. Morbidity varies in severity but commonly affects people in tropical and subtropical regions. It is estimated that over one billion people are infected by one or more species of parasitic roundworm [5], and the loss of disability-adjusted life years caused by these parasites ranks among the top of all Neglected Tropical Diseases [1]. In addition to their devastating impacts on human health, several parasitic nematode species infect a variety of crops and livestock. These infections cause severe economic burdens worldwide [6].

Parasitic nematodes are primarily treated using a limited number of anthelmintic drugs from one of the three major drug classes: benzimidazoles, nicotinic acetylcholine receptor agonists, and macrocyclic lactones (MLs). However, the efficacies of these anthelmintics can be limited by the ubiquitous resistance observed in veterinary parasites [7] and the emerging resistance in human parasites [8,9]. In many cases, resistance is highly heritable, suggesting the evolution of anthelmintic-resistant nematodes might occur under drug selection [10]. We must understand the drug mode of action and identify the genetic loci that contribute to resistance in parasitic nematodes to provide effective long-term treatments. Abamectin and ivermectin are two common MLs used to treat parasitic nematodes of agricultural, veterinary, or human importance [11]. However, widespread resistance to MLs has been reported and is a significant concern [10]. Genetic screens and selections performed in the laboratory-adapted reference strain of the free-living nematode *Caenorhabditis elegans* have identified three genes that encode glutamate-gated chloride (GluCl) channel subunits (*glc-1*, *avr-14*, and *avr-15*) and are targeted by MLs in *C. elegans* [12,13]. In contrast to *C. elegans* laboratory experiments, several resistant parasitic nematode isolates have been discovered that do not have mutations in genes that encode GluCl subunits [14,15], suggesting that alternative mechanisms of resistance to MLs must exist. Recently, quantitative trait loci (QTL) mappings in both free-living and parasitic nematode species have identified several genomic regions of interest containing genetic variation that confers drug resistance [14,16–20].

The identification of specific genes or variants involved in the molecular basis of drug resistance in parasitic nematodes can be challenging for several reasons. First, their life-cycles require hosts and are costly to maintain [21]. Second, most species do not have annotated genomes assembled into full chromosomes. To date, the most complete genome is *Haemonchus contortus*, which enables genetic mappings and comparative genomic approaches [14,22]. Finally, most species lack key molecular and genetic tools such as CRISPR-Cas9 genome editing [23]. By contrast, the free-living nematode *C. elegans* has a short life cycle that is easy to grow in the laboratory, a well annotated reference genome, and a plethora of molecular and genetic tools to characterize anthelmintic responses [19,20,24–26]. Genetic mappings using hundreds of genetically and phenotypically diverged wild strains of *C. elegans* collected around the world could identify novel loci that contribute to anthelmintic resistance across natural populations [20,24].

Here, we use genome-wide association and linkage mapping analyses to identify three large-effect QTL on chromosome V that contribute to abamectin resistance. One of these QTL was previously identified and is known to be caused by variation in the GluCl channel gene *glc-1* [24,26]. The remaining two QTL are novel. We used near-isogenic lines (NILs) to validate and narrow each QTL independently and identified promising candidate genes to test using CRISPR-Cas9 genome editing. We searched for orthologous genes within ivermectin-response QTL on chromosome V of *H. contortus* and discovered 40 genes present in QTL for both species. This study demonstrates the value of natural variation in the *C. elegans* population to identify candidate genes for resistance and enables understanding of the molecular mechanisms of anthelmintic resistance.

## Results

### Two different quantitative genetic mapping techniques reveal loci that underlie differential responses to abamectin

Anthelmintic resistance can be described as a function of nematode development and reproduction. To quantify *C. elegans* drug resistance, we previously developed a high-throughput assay that uses a flow-based device to measure the development and reproduction of thousands of animals across hundreds of independent strains (see Methods) [19,20,27–33]. In this assay, nematode development is described by a combination of animal length, animal optical density (body thickness and composition integrated by length), and normalized animal optical density (body thickness and composition normalized by length). Although length and optical density are often highly correlated, these three traits can each describe a unique aspect of development [31]. In addition to development, this assay describes nematode reproduction by an approximation of animal brood size. Using this assay, we exposed four genetically divergent strains (N2, CB4856, JU775, and DL238) to increasing doses of abamectin (**S1 File**). In the presence of abamectin, nematodes were generally smaller, less optically dense, and produced smaller broods compared to non-treated nematodes, suggesting an abamectin-induced developmental delay and decreased reproduction (**S1 Fig**). We also observed significant phenotypic variation among strains in response to abamectin, indicating that genetic variation might underlie differences in the abamectin response across the *C. elegans* species.

To investigate the genetic basis of natural abamectin resistance, we exposed 210 wild isolates to abamectin and measured their developmental rates and brood sizes (**S2 File**). These data were used to perform genome-wide association mappings that identified a total of six QTL across the four traits on chromosomes II, III, and V (**S2 Fig, S3 File**). The most significant QTL was detected for brood size on the right arm of chromosome V (VR) and was also detected for animal length (**Fig 1A**). Notably, this region (**Table 1**) includes a gene (*glc-1*) that encodes a subunit of a glutamate-gated chloride (GluCl) channel, which has been previously discovered to underlie phenotypic differences in both swimming paralysis [24] and survival [26] in the presence of abamectin. Additionally, a second QTL on the left arm of chromosome V (VL) underlies differences in both animal length and optical density (**Fig 1A**, **S2 Fig, S3 File**). This secondary region (**Table 1**) has not been identified in previous *C. elegans* QTL mapping studies [24,26] and thus can be considered a novel region underlying abamectin response.

**Fig 1.**
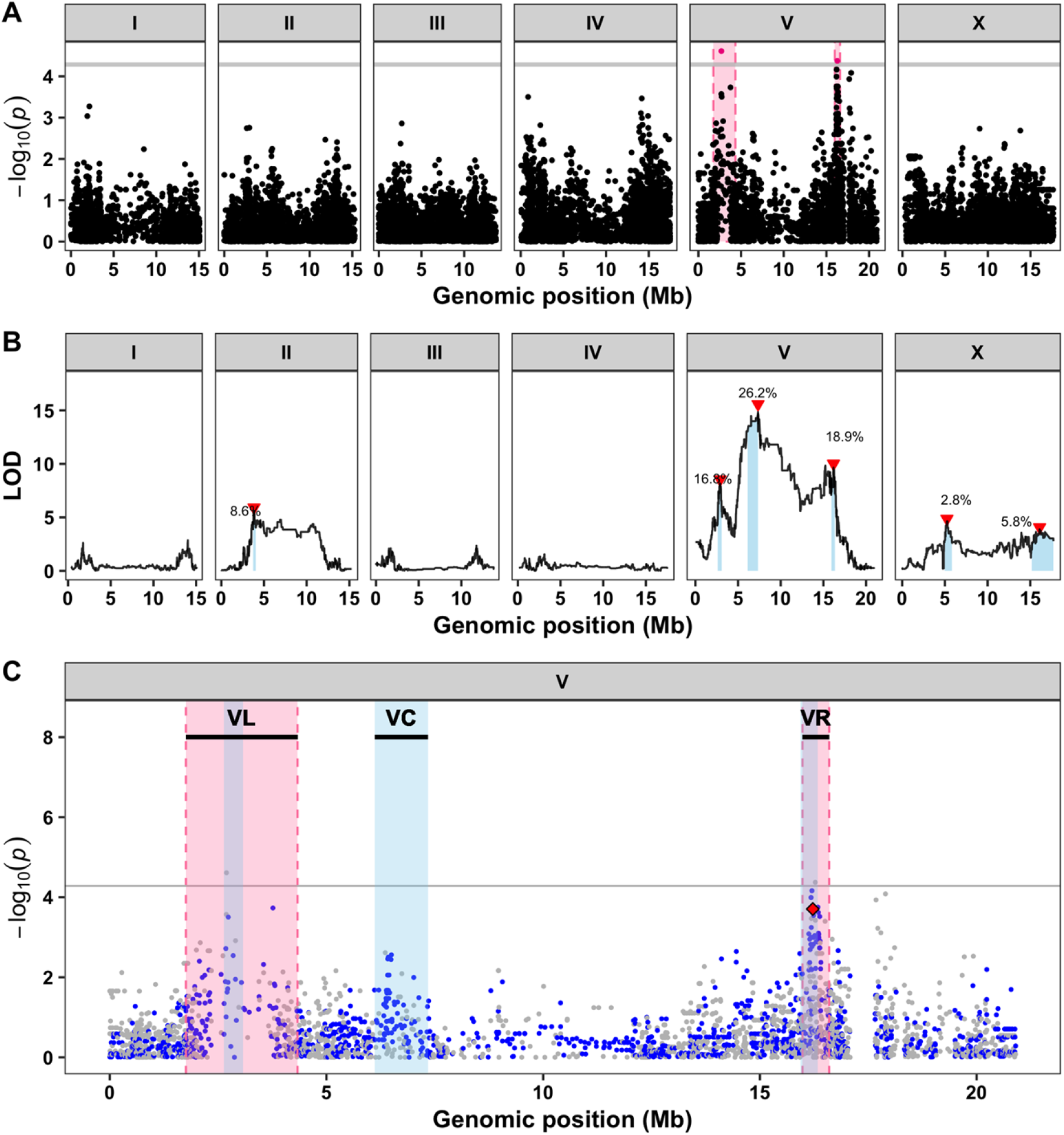
Three large-effect QTL on chromosome V control differences in abamectin responses. **A)** Genome-wide association mapping results for animal length (mean.TOF) are shown. Genomic position (x-axis) is plotted against the −log10(*p*) value (y-axis) for each SNV. SNVs are colored pink if they pass the genome-wide eigen-decomposition significance threshold designated by the grey line. The genomic regions of interest that pass the significance threshold are highlighted by pink rectangles. **B)** Linkage mapping results for optical density (mean.EXT) are shown. Genomic position (x-axis) is plotted against the logarithm of the odds (LOD) score (y-axis) for 13,003 genomic markers. Each significant QTL is indicated by a red triangle at the peak marker, and a blue rectangle covers the 95% confidence interval around the peak marker. The percentage of the total variance in the RIAIL population that can be explained by each QTL is shown above the QTL. **C)** Fine mapping of all common variants on chromosome V is shown. Genomic position (x-axis) is plotted against the −log10(*p*) values (y-axis) for each variant and colored by the genotype of the variant in the CB4856 strain (grey = N2 reference allele, blue = variation from the N2 reference allele). Genomic regions identified from linkage mapping analysis are highlighted in blue and genomic regions identified from association mapping are highlighted in pink. The horizontal grey line represents the genome-wide eigen-decomposition significance threshold. The red diamond represents the most significant variant in the gene *glc-1*.

**Table 1.** Genomic regions on chromosome V significantly correlated with abamectin resistance.

| QTL | Association mapping<sup>1</sup> | Linkage mapping<sup>2</sup> | NIL-defined interval |
| --- | --- | --- | --- |
| VL | V:1,757,246-4,333,001 | V:2,629,324-3,076,312 | V:1-3,120,167 |
| VC | NA | V:6,118,360-7,342,129 | V:5,260,997-5,906,132 |
| VR (*glc-1*)<sup>3</sup> | V:15,983,112-16,599,066 | V:15,933,659-16,336,743 | 13,678,801-19,303,558 |

<sup>1</sup>Mean animal length
<sup>2</sup>Mean animal optical density
<sup>3</sup>From previous data [24]

In parallel to the genome-wide association mapping, we measured animal development and reproduction in abamectin for a panel of 225 recombinant inbred advanced intercross lines (RIAILs) generated by a cross between the N2 and CB4856 strains [31] (**S4 File**). Linkage mapping analysis for all four traits identified a total of 14 QTL on chromosomes II, V, and X including three distinct QTL on chromosome V (**Fig 1B**, **S3 Fig, S5 File**). Two of the three QTL on chromosome V (VL and VR) overlap with the intervals identified using association mapping (**Fig 1C**, **Table 1**, **S4 Fig, S7 File**), suggesting that a single variant both present in the CB4856 strain and prevalent within the natural population might cause the differences in abamectin responses observed in both mapping populations. In fact, a four-amino-acid deletion in the gene *glc-1* that has been previously correlated with abamectin resistance is known to segregate not only among the *C. elegans* population but also between the N2 and CB4856 strains [24,26]. By contrast, the third locus on the center of chromosome V (VC) was only detected using linkage mapping, which suggests that a rare variant in the CB4856 strain underlies this QTL. Regardless, all three loci on chromosome V can be investigated further by leveraging the genetic variation between the N2 and CB4856 strains. At each of the three loci on chromosome V, RIAILs with the CB4856 allele were correlated with abamectin resistance (longer and denser animals) compared to RIAILs with the N2 allele (**S3 Fig, S4 and S5 Files**). To search for evidence of any genetic interactions between loci, we performed a two-dimensional genome scan. We found no significant interactions on chromosome V or otherwise (**S5 Fig, S6 File**), suggesting that each of the three loci additively contribute to abamectin resistance.

### Near-isogenic lines validate the independent abamectin-resistance loci on chromosome V

To empirically validate if genetic variation on chromosome V between the N2 and CB4856 strains contributes to abamectin resistance, we first generated chromosome substitution strains in which the entire chromosome V from the CB4856 strain was introgressed into the N2 genetic background or vice versa (**S8 and S9 Files**). We measured the development and reproduction of these strains and observed that the genotype on chromosome V significantly contributed to differences in abamectin resistance (**S10 File**). The strain with the CB4856 genotype on chromosome V introgressed into the N2 genetic background was significantly more resistant than the sensitive N2 strain (Tukey’s HSD, *p*-value = 2.47e-06) (**S6 Fig, S10 and S11 Files**). Similarly, the strain with the N2 chromosome V introgressed into the CB4856 genetic background was significantly more sensitive to abamectin compared to the resistant CB4856 strain (Tukey’s HSD, *p*-value = 0.0041, **S6 Fig, S10 and S11 Files**). These results confirm that genetic variation between the N2 and CB4856 strains at one or more loci on chromosome V contributes to the difference in abamectin resistance between these strains.

To demonstrate that genetic variation outside the *glc-1* locus contributes to the overall resistance phenotype observed in the chromosome substitution strains, we next generated a near-isogenic line (NIL) that contains the resistant CB4856 alleles at both the VL and VC loci and the sensitive N2 allele at the VR *glc-1* locus (**Fig 2A**, **S8 and S9 Files**). When tested, we observed that this NIL (ECA1059) was significantly more resistant to abamectin than the sensitive N2 strain (Tukey’s HSD, *p*-value = 8.83e-14) and less resistant than the resistant CB4856 strain (Tukey’s HSD, *p*-value = 1.57e-13, **Fig 2B**, **S11 and S12 Files**). This result indicates that genetic variation on the left and/or center of chromosome V contributes to the differences in abamectin resistance between the N2 and CB4856 strains.

**Fig 2.**
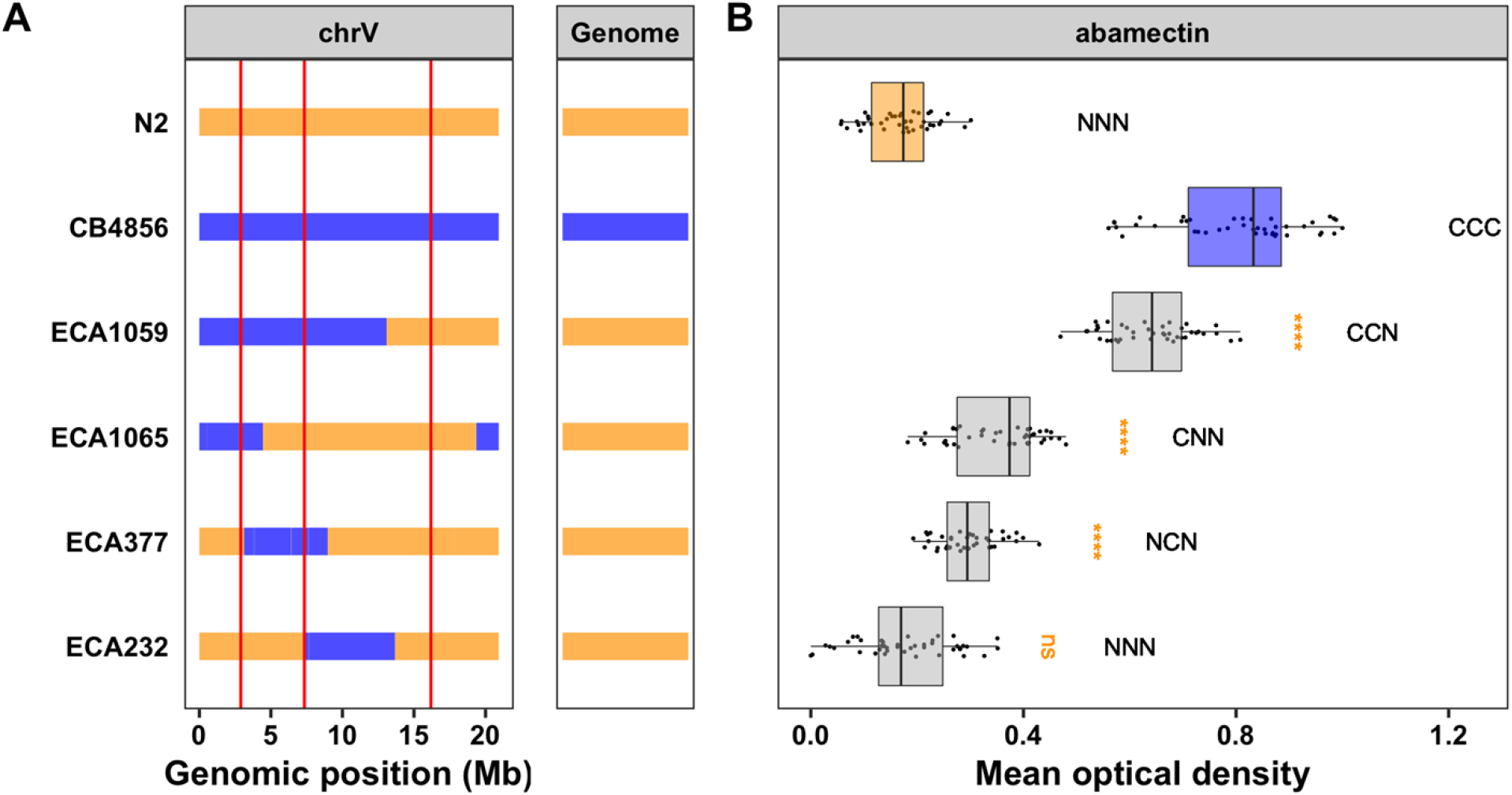
Near-isogenic lines confirm the additive effects of all three QTL. **A)** Strain genotypes are shown as colored rectangles (N2: orange, CB4856: blue) in detail for chromosome V (left box) and in general for the rest of the chromosomes (right box). The solid vertical lines represent the peak marker of each QTL. **B)** Normalized residual mean optical density in abamectin (mean.EXT, x-axis) is plotted as Tukey box plots against strain (y-axis). Statistical significance of each NIL compared to N2 calculated by Tukey’s HSD is shown above each strain (ns = non-significant (p-value > 0.05), *** = significant (p-values < 0.0001). Predicted genotypes at the three QTL are shown above each strain from VL to VC to VR (N = N2 allele, C = CB4856 allele).

To further isolate the VL and VC QTL independently, we generated three NILs that each contain approximately 5 Mb of the CB4856 genome introgressed into the N2 genetic background at different locations on chromosome V so that they tile across the introgressed region in ECA1059 (**Fig 2A**, **S8 and S9 Files**). The strain ECA232 was not significantly more resistant to abamectin compared to the N2 strain (Tukey’s HSD, *p*-value = 0.997), suggesting that this NIL has the sensitive N2 alleles at all three loci on chromosome V (**Fig 2B**, **S11 and S12 Files**). Alternatively, both ECA1065 and ECA377 were significantly more resistant to abamectin than the N2 strain (Tukey’s HSD, *p*-values = 1.45e-13 and 5.44e-09, respectively), suggesting that the introgressed regions in both of these NILs contain one or more resistant loci (**Fig 2B**, **S11 and S12 Files**). Furthermore, because both strains are less resistant than the NIL with two CB4856 alleles (ECA1059-ECA1065 *p*-value = 8.83e-14 (Tukey’s HSD), ECA1059-ECA377 *p*-value = 8.83e-14 (Tukey’s HSD)), we can deduce that ECA1065 and ECA377 each contain one resistant locus (**Fig 2B**, **S11 and S12 Files**). The introgressions in these two NILs overlap by 1.3 Mb, which leaves two possibilities: either this overlapped region (V:3,120,168-4,446,729) contains a single QTL shared by the two NILs or each NIL validates a separate QTL within the non-overlapping introgressed regions. Because we identified three QTL from linkage mapping, we believe the latter case in which ECA1065 has the CB4856 allele for the VL locus and ECA377 has the CB4856 allele for the VC locus (**Table 1**, **S7 Fig**).

The VL QTL is contained within the CB4856 introgression in the NIL ECA1065 and is defined by the CB4856 introgression in the NIL ECA377 (V:1-3,120,167). This NIL-defined interval overlaps with the genomic regions identified from both linkage and association mapping (**S7 Fig**, **Table 1**). Using the smallest of these overlapping regions (V:2,629,324-3,076,312), we identified a total of 164 genes in the interval. A change in phenotype is most commonly driven by either genetic variation that alters the amino acid sequence of a protein (protein-coding genetic variation) or genetic variation that affects expression of one or more genes (expression variation). Using previously published gene expression data [32,34] and genetic variant data accessed from the *C. elegans* Natural Diversity Resource (CeNDR; elegansvariation.org), we eliminated 61 candidate genes because 26 genes did not harbor any genetic variation between the N2 and CB4856 strains and 35 genes had no protein-coding or expression variation. The remaining 103 candidate genes had either protein-coding or expression variation linked to this region (**S13 File**). Although none of these 103 genes encode a GluCl channel, three genes with variation in expression between the N2 and CB4856 strains do encode a cytochrome P450 (*cyp-35C1*, *cyp-33E3*, and *cyp-33E2*) and one encodes a UDP-glycosyltransferase (*ugt-53*). Previous studies have shown an upregulation of such metabolic genes in response to benzimidazole treatment [35]. Therefore, it is possible that differential expression of key metabolic genes between the N2 and CB4856 strains causes variation in abamectin resistance.

To further test the role of gene expression variation in the abamectin resistance phenotype, we measured animal development and reproduction in abamectin for 107 RIAILs for which we had existing gene expression data [32,34] (**S14 File**) and performed mediation analysis for each of the 28 genes with an eQTL in the region (**S8 Fig, S15 File**). The top three candidates from this analysis included a gene with unknown function (*F54E2.1*), an NADH Ubiquinone Oxidoreductase (*nuo-5*), and *ugt-53*. The gene *nuo-5* is a NADH-ubiquinone oxidoreductase and part of the mitochondrial complex I [36,37]. Animals deficient for *nuo-5* are more sensitive to the anthelmintic levamisole, and have defective cholinergic synaptic function [37]. Interestingly, the CB4856 strain has lower expression of both *nuo-5* and *ugt-53*, which suggests that if either of these genes are causal it is likely through an indirect mechanism. Regardless, we have previously shown that mediation analysis is a strong tool for predicting candidate genes with expression variation [32,38], and this analysis provides more evidence for the potential role of *ugt-53* or *nuo-5* in the abamectin response. However, it is important to note that this genomic interval also lies within a hyper-divergent region of the genome marked by extremely high levels of structural and single-nucleotide variation in the CB4856 strain as well as many other strains [39]. This extreme level of genetic variation could, in some cases, even cause a different composition of genes than found in the N2 reference strain. Because candidate gene predictions could be highly impacted by this divergent region, we turned our focus to the VC QTL.

The VC QTL is contained within the CB4856 introgression in the NIL ECA377 and is defined by the CB4856 introgressions in the NILs ECA1065 and ECA232 (V:4,446,729-7,374,928) (**Fig 2**, **S7 Fig**). This large 3 Mb interval overlaps with the genomic region identified using linkage mapping (**S7 Fig**, **Table 1**). We next attempted to narrow this region further by generating additional NILs with smaller introgressions (**Fig 3A, S8 and S9 Files**) and measuring the abamectin resistance of these strains. All four NILs with the CB4856 introgression on the center of chromosome V were significantly more resistant (as defined by nematode length) than the N2 strain (Tukey’s HSD, *p*-values < 2.83e-06, **Fig 3**, **S11 and S16 Files**). The results for optical density were similar, but the variation within strains was higher (**S9 Fig, S11 and S16 Files**). These results suggest that the QTL position is contained within the smallest introgression, ECA632 (V:5,260,997-5,906,132) (**Table 1, S7 Fig**). Interestingly, this NIL-defined genomic region does not overlap with the confidence interval identified using linkage mapping (**Table 1**, **S7 Fig**). Regardless, we prioritized 103 potential candidate genes with either protein-coding variation and/or variation in gene expression linked to this 645 kb region (**S13 File**). Notably, the glutamate-gated chloride channel, *glc-3*, resides within this narrowed region (V:5,449,287, **S7 Fig**). GLC-3 was shown to be activated by ivermectin when heterologously expressed in *Xenopus laevis oocytes* [40]. However, it has yet to be identified in mutant screens nor directly tested for ML resistance in *C. elegans*. The CB4856 strain has three rare variants in *glc-3*, including a single missense variant (I439F). It is possible that one or more of these variants causes increased resistance to abamectin in the CB4856 strain. In addition to *glc-3*, this list of prioritized candidate genes includes one cytochrome P450 (*cyp-35B2*) and two UDP-glycosyltransferases (*ugt-51* and *H23N18.4*). Mediation analysis for the 48 genes with an eQTL that maps to this region highlighted several potential candidates, most noticeably *H23N18.4* (**S8 Fig, S15 File**). Variation in one or more of these genes might contribute to differences in abamectin resistance between the N2 and CB4856 strains.

**Fig 3.**
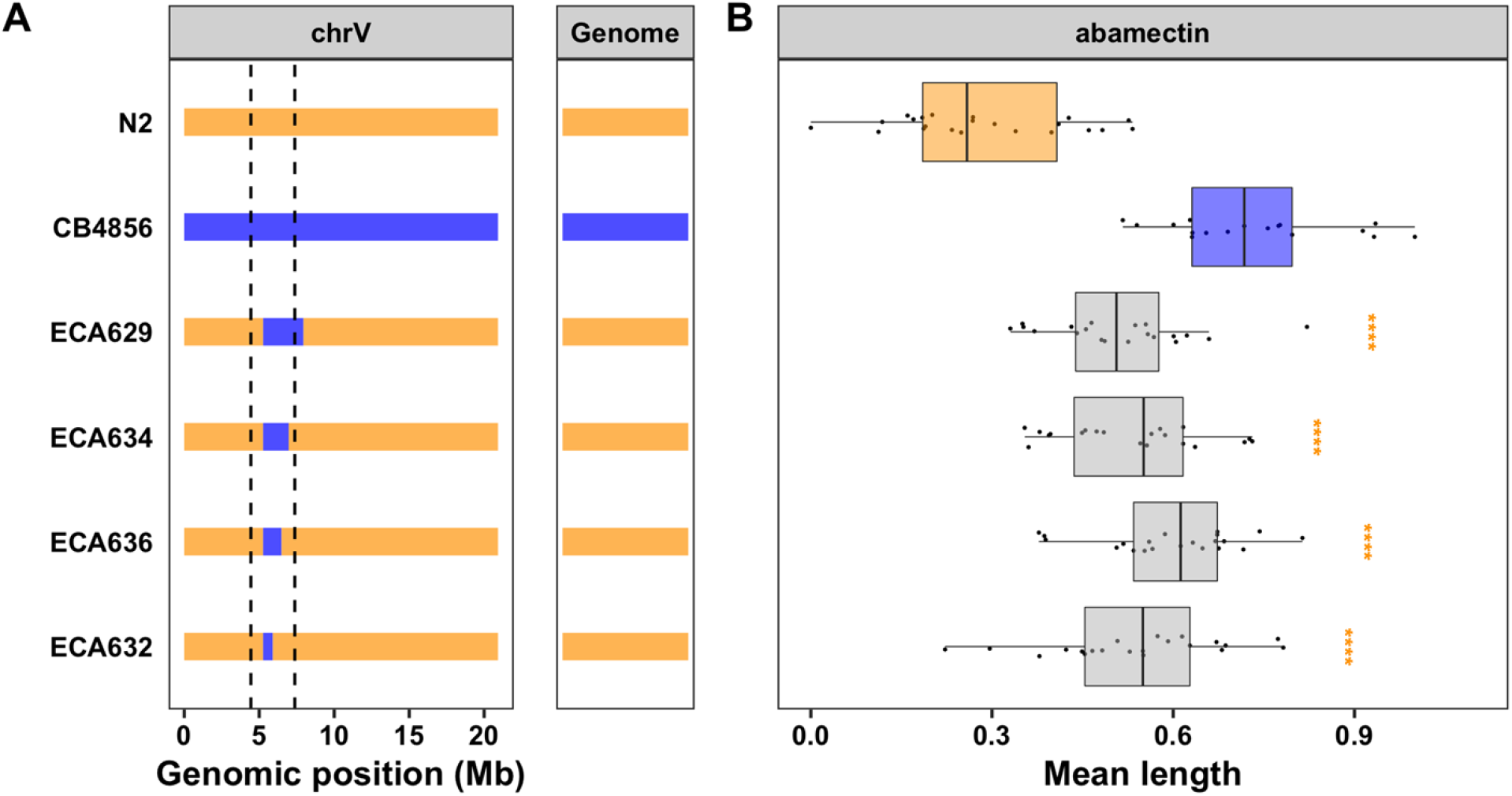
NILs isolate and narrow the VC QTL. **A)** Strain genotypes are shown as colored rectangles (N2: orange, CB4856: blue) in detail for chromosome V (left) and in general for the rest of the chromosomes (right). The dashed vertical lines represent the previous NIL-defined QTL interval for VC. **B)** Normalized residual mean animal length in abamectin (mean.TOF, x-axis) is plotted as Tukey box plots against strain (y-axis). Statistical significance of each NIL compared to N2 calculated by Tukey’s HSD is shown above each strain (ns = non-significant (p-value > 0.05), *** = significant (p-values < 0.0001).

### The candidate gene *lgc-54* likely does not cause macrocyclic lactone resistance

Several previous studies in parasitic nematode species have identified QTL and candidate genes that might underlie responses to ivermectin, an ML closely related to abamectin. Introgression mapping in the sheep parasite *Teladorsagia circumcincta* identified several candidate genes, including the ortholog of the *C. elegans* gene *lgc-54* [16], but this genome is highly fragmented and genomic locations are likely not correct. Regardless, this gene, similar to *glc-1*, encodes a ligand-gated chloride channel, but it has yet to be directly implicated in ML resistance. Interestingly, *lgc-54* is on the center of the *C. elegans* chromosome V (6.8 Mb) within the confidence interval for the VC QTL defined by the linkage mapping experiment but outside the NIL-defined interval (**Table 1**, **S7 Fig**). Furthermore, the CB4856 strain harbors several genetic variants in this gene, including a single missense variant (H42R) unique to the CB4856 strain. To test if *lgc-54* plays a role in abamectin resistance in *C. elegans*, we exposed two independent *lgc-54* mutants to abamectin and measured animal development and reproduction. Both *lgc-54* mutants were significantly more resistant than the N2 strain (Tukey’s HSD, *p*-values < 0.0004), which would normally provide evidence for the role of *lgc-54* in abamectin resistance (**Fig 4A**, **S11 and S17 Files**). However, we noticed that these mutants grew much slower in the control conditions than both the N2 and CB4856 strains (**Fig 4B**, **S17 File**). This growth defect suggests that the observed resistance is likely an artifact of the statistical regression analysis, suggesting that *lgc-54* does not play a role in abamectin resistance. Regardless, this system provides a strong platform for experimentally validating putative resistance alleles from parasitic nematodes.

**Fig 4.**
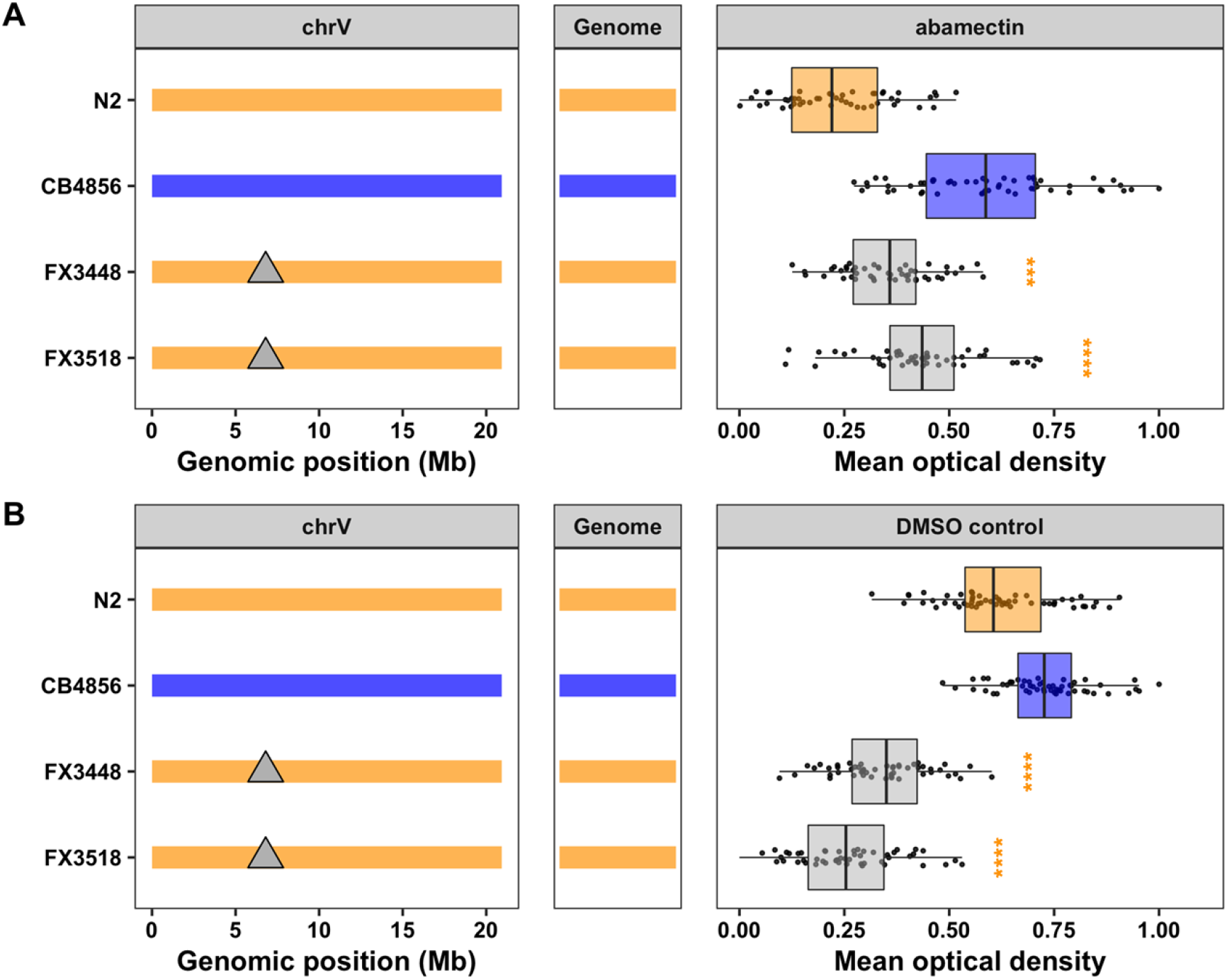
Testing the role of *lgc-54* in the *C. elegans* abamectin response. Strain genotypes are shown as colored rectangles (N2: orange, CB4856: blue) in detail for chromosome V (left) and in general for the rest of the chromosomes (center). Grey triangles represent mutations in the *lgc-54* gene. On the right, normalized residual mean optical density in abamectin (mean.EXT, x-axis) **(A)** or normalized mean optical density in DMSO control **(B)** is plotted as Tukey box plots against strain (y-axis). Statistical significance of each deletion strain compared to N2 calculated by Tukey’s HSD is shown above each strain (ns = non-significant (p-value > 0.05), **** = significant (p-value < 0.0001).

### Overlap of *C. elegans* and parasitic nematode candidate genes for macrocyclic lactone resistance

Recently, two large-effect QTL on chromosome V (37-42 Mb and 45-48 Mb) were identified in response to ivermectin treatment in *H. contortus*, a gastrointestinal parasite of small ruminants [14]. The authors found that these regions did not contain any orthologs of candidate ML resistance genes identified in *C. elegans* previously. Because the QTL for MLs in both species are found on chromosome V and the chromosomal gene content is conserved between *C. elegans* and *H. contortus* [22], orthologs present in the QTL of both species can suggest conserved resistance mechanisms. To compare the genes in the newly defined QTL (VL and VC) to those ivermectin QTL defined for *H. contortus*, we identified one-to-one orthologs across the two species and compared chromosomal positions (**Fig 5**, **S18 File**). For this analysis, larger and more conservative genomic regions were selected to represent the *C. elegans* QTL (V:1-3,120,167 and V:5,260,997-7,342,129). Consistent with previous work that found that linkage groups are highly conserved but synteny (or gene order) is not [14], we found that the genes within the *C. elegans* QTL are distributed throughout the *H. contortus* chromosome V, with only 40 (21.62%) of the 185 one-to-one orthologs present in one of the two *H. contortus* QTL. Regardless, we investigated functional annotations of the one-to-one orthologs shared between the QTL and found that none have annotations that have previously been associated with ML resistance.

**Fig 5.**
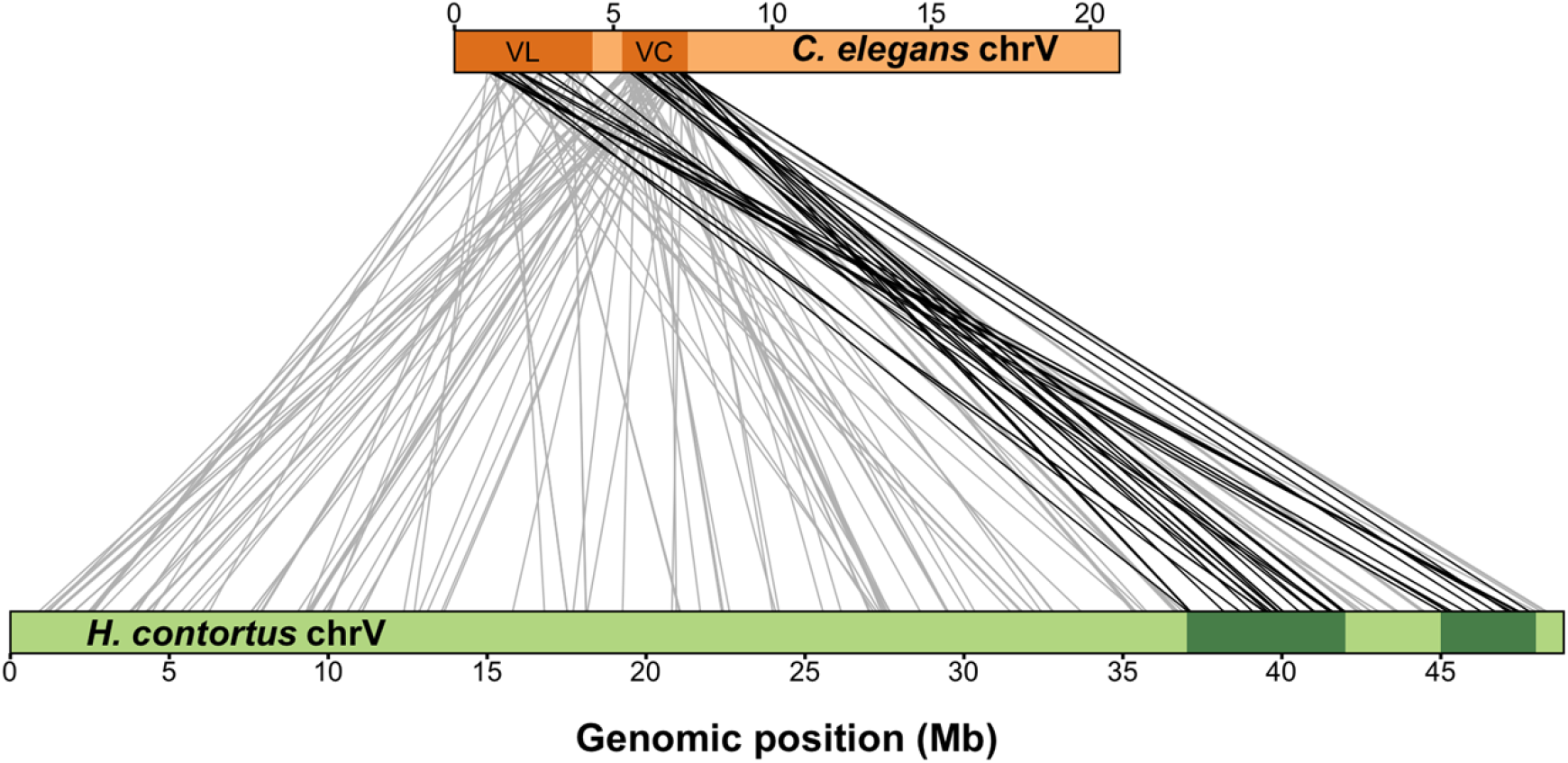
Synteny between major-effect *C. elegans* abamectin QTL and *H. contortus* ivermectin QTL on chromosome V. Synteny plot showing orthologous genes between *C. elegans* (top) and *H. contortus* (bottom) chromosome V. Genomic regions of interest are highlighted with dark orange (*C. elegans*) or dark green (*H. contortus*) rectangles, and orthologous gene pairs are represented by connecting lines. Only genes that lie within either the VL or VC *C. elegans* QTL with a one-to-one ortholog on *H. contortus* chromosome V are shown (185 pairs). Grey lines represent a gene pair where the *H. contortus* ortholog is not within a region of interest (145 pairs), and black lines represent a gene pair where the *H. contortus* ortholog is within a region of interest (40 pairs).

However, genes associated with anthelmintic resistance can be members of large gene families, like UDP-glycosyltransferases (UGTs) or cytochrome P450 enzymes (CYPs), and often do not have one-to-one orthologs. For this reason, we searched for orthogroups with more than one gene in one or both species within both QTL (**S19 File**). The complexity of comparing gene families between *C. elegans* and *H. contortus* prohibits searches for each of the 1169 genes in the VL and VC regions. Therefore, we narrowed our comparison to include one-to-many, many-to-one, and many-to-many orthologs of previously described *C. elegans* candidate genes. This approach identified orthologs for the CYP and UGT families in the *C. elegans* QTL, but none were present in the *H. contortus* QTL. Additionally, we investigated two candidate genes, *glc-3* and *nuo-5*, that we identified in this study. The glutamate-gated chloride channel subunit gene, *glc-3*, is located in the VC region. It has a one-to-one ortholog on *H. contortus* chromosome V, but this ortholog, *HCON_00148840*, is located to the left of the *H. contortus* QTL region (27.6 Mb). However, the QTL location and confidence intervals depend on the underlying statistics and studied populations, so this candidate gene could still play a role in *H. contortus.* Alternatively, the one-to-one ortholog of the NADH ubiquinone oxidoreductase gene, *nuo-5*, is found on the far left of *H. contortus* chromosome V (2.47 Mb) and is unlikely to underlie the *H. contortus* QTL. Overall, our analysis indicates some orthologs are shared between QTL or neighboring regions from the two species. These orthologs should be prioritized for further studies of ML resistance.

## Discussion

In this study, we used *C. elegans* genome-wide association and linkage mapping analyses to identify three QTL on chromosome V that influence responses to the ML abamectin. One of these QTL overlaps with the previously identified GluCl gene *glc-1* [24,26]. However, the remaining two QTL are novel and might overlap with ivermectin QTL from parasitic nematodes, showing the power of using *C. elegans* to discover conserved ML resistance genes. We used NILs to validate and narrow each QTL independently. Additionally, we compared genes in our QTL with genes in two ivermectin QTL for *H. contortus* [14] and identified 40 shared orthologs. However, none of these 40 genes were strong candidates based on previous implications in ML resistance. Although we were unable to discover the specific causal genes or variants, we suggest several candidate genes within the narrowed genomic intervals that could play roles in ML resistance.

### Different mapping populations and techniques detect both similar and distinct QTL

Abamectin resistance has now been mapped in *C. elegans* using three different experimental mapping techniques, five mapping populations, and several distinct traits [24,26]. Common to all these studies is the large-effect QTL detected on the right arm of chromosome V (VR locus), likely controlled by natural genetic variation in the gene *glc-1*. Here, we validated that genetic variation on the right arm of chromosome V contributes to abamectin resistance using NILs. However, further validation of the specific variants that affect the function of *glc-1* in ML response is still needed. Regardless, the overlap of QTL between these three studies and across mapping populations suggest that the causal variant is commonly occuring and has a strong effect on ML resistance in *C. elegans* across a variety of traits.

Unlike the previous two *C. elegans* abamectin resistance mappings, we identified several additional QTL, most significantly two novel QTL on chromosome V (VL and VC). Because the previous studies that only detected *glc-1* measured swimming paralysis and survival in abamectin, it is possible that the VL and VC novel QTL could underlie differences in nematode development as measured in the high-throughput growth and fecundity assay performed here. These differences in traits and underlying QTL could suggest a complex nematode response to MLs. Although the GluCl-encoding gene, *glc-1*, underlies many of the differences across *C. elegans* strains, we show that these other loci underlie differences in development and physiology independent of *glc-1*.

The VL QTL was detected using both linkage and association mapping. The overlap of QTL from these two distinct mapping methods suggests that, like *glc-1*, genetic variation in the CB4856 strain is also found commonly across the species. Alternatively, the VC QTL was detected only with linkage mapping analysis, suggesting that either the causal variant in the CB4856 strain is rare across the species or does not cause a resistant phenotype in other wild strains. We used near-isogenic lines (NILs) to empirically validate both QTL independently. Interestingly, we show that a 645 kb region located outside of the statistically defined linkage mapping confidence interval is sufficient to cause resistance in an otherwise sensitive genetic background. Although several reasons could account for this discrepancy, it is likely that several small-effect loci on the center of chromosome V each contribute to the overall abamectin-resistance phenotype observed in the CB4856 strain. In the recombinant population, all of these loci are jointly evaluated causing the QTL to be defined at a specific location. However, NILs are able to isolate small regions of the genome and these multiple effects can now be detected independently [38,41]. This study emphasizes that, although powerful, QTL mapping is ultimately a statistical method that can be influenced by experimental differences and that it is essential to validate QTL before drawing conclusions about the genomic location of the causal variant. Although validating QTL can be difficult in many parasitic nematode species, our ability to validate QTL in *C. elegans* is a strength of this model organism, emphasizing the need for the parasitic nematode and *C. elegans* communities to work together to push forward a cycle of discovery to understand anthelmintic modes of action and mechanisms of resistance [23].

### Shared niches provide the same selective pressures for soil transmitted helminths and *C. elegans*

The potential overlap of QTL for ML resistance between *C. elegans* and parasitic nematodes suggests that the loci that confer resistance to MLs could be conserved across several nematode species. It is believed that parasitic nematodes are resistant to anthelmintic drugs because high levels of standing genetic variation harbor existing resistance alleles in a population [42,43]. Soil transmitted helminths such as *H. contortus*, spend part of their life cycle in soil or rotting vegetation, an environment that overlaps with the niche associated with the free-living *C. elegans* [44,45]. Selective pressures in this environment could originate from natural toxic compounds produced by soil-dwelling bacteria and fungi from which many anthelmintic drugs are derived [46–48]. MLs, like abamectin and ivermectin, are fermentation products of *Streptomyces avermitilis* [49]. This gram-positive bacteria was originally isolated in Japan, but shown to grow in a variety of substrates [50] and its presence in the soil could select for naturally resistant nematodes. Additionally, synthetic anthelmintic compounds are common soil and water pollutants in some areas and can be found in runoff from farms that use anthelmintics to treat agriculture or livestock [51,52]. This exposure to the same selective pressures and the known genetic diversity in the *C. elegans* species suggests a method for how free-living nematodes might acquire variation in the same resistance genes or gene families as parasitic nematodes. In one example, it was shown that recent selective pressures have likely acted on the *C. elegans ben-1* locus, causing many novel putative loss-of-function alleles across the population despite evolutionary constraint on *ben-1* beta-tubulin function [20]. This conclusion again highlights the relevance of using the experimentally tractable *C. elegans* as a model to study anthelmintic resistance in parasitic nematode species.

### Overlap of chromosome V macrocyclic lactone resistance loci between *C. elegans* and *H. contortus*

Genome-wide analyses in both *C. elegans* and *H. contortus* identified chromosome V QTL. Because gene contents on chromosomes are highly conserved, we compared these QTL intervals to look for conserved candidate genes. We initially focused on one-to-one orthologs between the VL and VC QTL in *C. elegans* and both QTL in *H. contortus* and found 40 genes but none were obvious new candidates for ML resistance. This finding does not eliminate the possibility that similar responses underlie the QTL in both species. The VL and VC QTL contain several gene families, including *ugt* and *cyp* genes, which have been implicated in resistance previously [53,54]. Gene families often evolve rapidly in response to the environment and are good candidates to confer resistance [55]. Because comparisons of all gene families or ortholog groups in the VL and VC QTL is difficult, we focused on ortholog groups for a subset of previously described candidate gene families (*ugt* and *cyp*). Neither of these families had orthologs in QTL for both species.

The absence of shared candidates between the QTL could be explained by the statistical nature of the mapping approaches. The glutamate-gated chloride channel subunit gene, *glc-3*, is located in the *C. elegans* VC QTL. Although the *H. contortus* ortholog *HCON_00148840* is not located in one of the QTL, it is found on chromosome V to the left of the defined QTL region (27.6 Mb) and could underlie that QTL. Alternatively, *HCON_00148840* could be a candidate gene but not vary in the *H. contortus* isolates used in the study [14]. Genome-wide association studies correlate genotype with phenotype. These studies depend on genetic variation in the tested strains and do not provide conclusive evidence on causal connections of candidate genes to resistance. Further studies should use additional *H. contortus* isolates to map genomic regions that correlate with ML resistance. The role of candidate genes such as *glc-3* should be tested in *C. elegans*.

When studies in *C. elegans* and *H. contortus* do point to the same candidate genes, variants that confer resistance in parasitic nematode species can be more easily validated in the experimentally tractable *C. elegans* model. We showed this approach by testing the *T. circumcincta* candidate gene (*lgc-54*) for ML resistance in *C. elegans*. Our results suggest that loss of *lgc-54* does not cause increased resistance to abamectin in *C. elegans*. A possible caveat of this study is that the decreased fitness of these mutants make the results more difficult to interpret. However, the decreased fitness could also be indicative of an essential role of *lgc-54* in nematode fitness. This study demonstrates the power of functional validation in model systems like *C. elegans* to experimentally test hypotheses for candidate genes with one-to-one orthologs. To study the resistance conferred by genes that have multiple orthologs in one or both species, genome-editing could be used to make *C. elegans* gene content resemble parasitic nematodes [23,56]. Additionally, CeNDR can be used to identify strains that have similar gene contents as found in *H. contortus* [39,57].

### Power of QTL mapping in *C. elegans* to identify causal genes underlying anthelmintic resistance

This study highlights the benefits of communication between the parasite and *C. elegans* communities. Genetic mappings, screens, and selections are more easily performed in free-living nematodes and ultimately discover drug targets and mechanisms of action. However, it is important that these findings are then translated back to parasitic nematodes to confirm that genes found in *C. elegans* are responsible for drug resistance in parasites. Alternatively, candidate genes can be identified from parasitic nematode field samples and direct *C. elegans* studies to test specific genes in anthelmintic resistance traits. The potential overlap between QTL for ML resistance in *C. elegans* and *H. contortus* [14] strengthens this approach and suggests that the variants in our mapping population might also confer resistance in *H. contortus* and perhaps other parasitic nematode species. Future studies to discover the causal genes and variants underlying our two novel QTL (VL and VC) could be informative to parasitologists and help treat infected individuals more effectively.

## Materials and Methods

### Strains

Animals were grown on modified nematode growth media (NGMA) containing 1% agar and 0.7% agarose at 20°C and fed the *E. coli* strain OP50 [58]. A total of 225 recombinant inbred advanced intercross lines (RIAILs) were assayed for QTL mapping (set 2 RIAILs) [31]. These RIAILs were derived from a cross between QX1430, which is a derivative of the canonical laboratory N2 strain that contains the CB4856 allele at the *npr-1* locus and a transposon insertion in *peel-1*, and a wild isolate from Hawaii (CB4856). A second set of 107 RIAILs generated previously between N2 and CB4856 [34] (set 1 RIAILs) were phenotyped for mediation analysis. Near-isogenic lines (NILs) were generated previously by backcrossing a RIAIL of interest to either the N2 or CB4856 strain for several generations using PCR amplicons covering insertion-deletion (indels) variants to track the introgressed region [28]. NILs were whole-genome sequenced to verify that only the targeted introgressed regions had been crossed. The *lgc-54* mutant strains (FX3448 and FX3518) were obtained from the National BioResource Project (Japan). All NILs and resources used to generate NILs are listed in the Supplementary Material. Strains are available upon request or from the *C. elegans* Natural Diversity Resource (CeNDR, elegansvariation.org) [57].

### High-throughput fitness assays

A high-throughput fitness assay described previously [31] was used for all phenotyping assays. In summary, each strain was passaged and amplified on NGMA plates for four generations, bleach-synchronized, and 25-50 embryos were aliquoted into 96-well microtiter plates at a final volume of 50 μL K medium [59]. After 12 hours, arrested L1s were fed HB101 bacterial lysate (Pennsylvania State University Shared Fermentation Facility, State College, PA; [60]) at a final concentration of 5 mg/mL in K medium. Animals were grown for 48 hours at 20°C with constant shaking. Three L4 larvae were then sorted into new 96-well microtiter plates containing 10 mg/mL HB101 bacterial lysate, 50 μM kanamycin, and either 1% DMSO or abamectin dissolved in 1% DMSO using a large-particle flow cytometer (COPAS BIOSORT, Union Biometrica; Holliston, MA). Sorted animals were grown for 96 hours at 20°C with constant shaking. The next generation of animals and the parents were treated with sodium azide (50 mM in 1X M9) to straighten their bodies for more accurate length measurements. Animal length (mean.TOF), optical density integrated over animal length (mean.EXT), and brood size (norm.n) were quantified for each well using the COPAS BIOSORT. Nematodes get longer (animal length) and become thicker and more complex (optical density) over developmental time, and these two traits are correlated with one another. Because these two traits are highly correlated, we also generated a fourth trait (mean.norm.EXT) that normalizes the optical density by length (EXT/TOF) in order to provide a means to compare optical densities regardless of animal lengths.

Phenotypic measurements collected by the BIOSORT were processed and analyzed using the R package *easysorter* [61] as described previously [28]. Differences among strains within the control conditions were controlled by subtracting the mean control-condition value from each drug-condition replicate for each strain using a linear model (*drug_phenotype ~ mean_control_phenotype)*. In this way, we are addressing only the differences among strains that were caused by the drug condition and the variance in the control condition does not affect the variance in the drug condition. An R shiny web app was previously developed [38] to visualize the results from the high-throughput assays and can be found here: https://andersen-lab.shinyapps.io/NIL_genopheno/.

### Abamectin dose response

Four genetically divergent strains (N2, CB4856, JU775, and DL238) were treated with increasing concentrations of abamectin using the standard high-throughput assay described above. A concentration of 5 μM abamectin (Sigma, #31732-100MG) in DMSO was selected for the linkage mapping experiments and 7.5 nM abamectin in DMSO was selected for the genome-wide association mapping and NIL experiments. These concentrations provided a reproducible abamectin-specific effect that maximizes between-strain variation and minimizes within-strain variation across the three traits. The higher concentration in the linkage mapping experiment falls into the range of previously reported *in vitro* assays, and the lower concentration in the GWA assay was meant to capture a wider range of responses found in the natural population.

### Genome-wide association mapping

A total of 210 wild isolates were phenotyped in both abamectin and DMSO using the standard high-throughput assay described above. A genome-wide association mapping was performed for animal optical density (mean.EXT), normalized optical density (mean.norm.EXT), length (mean.TOF), and brood size (norm.n) using the R package *cegwas2* (https://github.com/AndersenLab/cegwas2-nf) as described previously [27,33]. Genotype data were acquired from the latest VCF release (release 20200815) from CeNDR. We used BCFtools [62] to filter variants below a 5% minor allele frequency and variants with missing genotypes and used PLINK v1.9 [63,64] to prune genotypes using linkage disequilibrium. The additive kinship matrix was generated from the 21,342 markers using the *A.mat* function in the *rrBLUP* R package [65]. Because these markers have high LD, we performed eigen decomposition of the correlation matrix of the genotype matrix to identify 963 independent tests [27]. We performed genome-wide association mapping using the *GWAS* function from the *rrBLUP* package. Significance was determined by an eigenvalue threshold set by the number of independent tests in the genotype matrix [27]. Confidence intervals were defined as +/− 150 SNVs from the rightmost and leftmost markers that passed the significance threshold.

### Linkage mapping

A total of 225 RIAILs [31] were phenotyped in abamectin and DMSO using the HTA described above. Linkage mapping was performed on the measured traits using the R package *linkagemapping* (https://github.com/AndersenLab/linkagemapping) as described previously [28,32]. The cross object derived from the whole-genome sequencing of the RIAILs containing 13,003 SNVs was merged with the RIAIL phenotypes using the *merge_pheno* function with the argument *set = 2*. A forward search (*fsearch* function) adapted from the *R/qtl* package [66] was used to calculate the logarithm of the odds (LOD) scores for each genetic marker and each trait as *−n(ln(1-R^2^)/2ln(10))* where R is the Pearson correlation coefficient between the RIAIL genotypes at the marker and trait phenotypes [67]. A 5% genome-wide error rate was calculated by permuting the RIAIL phenotypes 1000 times. QTL were identified as the genetic marker with the highest LOD score above the significance threshold. This marker was then integrated into the model as a cofactor and mapping was repeated iteratively until no further QTL were identified. Finally, the *annotate_lods* function was used to calculate the effect size of each QTL and determine 95% confidence intervals defined by a 1.5 LOD drop from the peak marker using the argument *cutoff* = *“proximal”*.

### Mediation analysis

107 RIAILs (set 1 RIAILs) were phenotyped in both abamectin and DMSO using the high-throughput assay described above. Microarray expression for 14,107 probes were previously collected from the set 1 RIAILs [68], filtered [58], and mapped using linkage mapping with 13,003 SNPs [32]. Mediation scores were calculated with bootstrapping using *calc_mediation* function from the *linkagemapping* R package which uses the *mediate* function from the *mediation* R package [69] as previously described [32] for each of the probes with an eQTL within a genomic region of interest. Briefly, a mediator model (expression ~ genotype) and an outcome model (phenotype ~ expression + genotype) were used to calculate the proportion of the QTL effect that can be explained by variation in gene expression. All expression and eQTL data are accessible from the *linkagemapping* R package.

### Comparing macrocyclic lactone QTL between *C. elegans* and *H. contortus*

The *C. elegans* (WS273) and *H. contortus* (PRJEB506, WBPS15) [22] protein and GFF3 files were downloaded from WormBase [70] and WormBase ParaSite [71], respectively. The longest isoform of each protein-coding gene was selected using the agat_sp_keep_longest_isoform.pl script from AGAT (version 0.4.0) [72]. Filtered protein files were clustered into orthologous groups (OGs) using OrthoFinder (version 2.4.0; using the parameter *-og*) [73] and one-to-one OGs were selected. A Python script (available at https://github.com/AndersenLab/abamectin_manuscript) was used to collect the coordinates of all *C. elegans* genes within one of the identified QTL along with the coordinates of the corresponding *H. contortus* orthologs. These coordinates were used to compare synteny between the *C. elegans* abamectin QTL defined here and the *H. contortus* ivermectin QTL defined previously [14] (V: 37-42 Mb and V: 45-48 Mb).

### Statistical analysis

All gene position data for the *C. elegans* genome were collected using WormBase WS273. For NIL assays, complete pairwise strain comparisons were performed on drug residual phenotypes using a *TukeyHSD* function [74] on an ANOVA model with the formula *phenotype* ~ *strain*. All data and scripts to generate figures can be found at https://github.com/AndersenLab/abamectin_manuscript.

## Acknowledgements

We would like to thank members of the Andersen laboratory for their helpful comments on the manuscript. We would also like to thank WormBase for hosting both parasite and *C. elegans* genomes and annotations and the *C. elegans* Natural Diversity Resource (NSF CSBR 1930382) for data and tools critical for our analysis. We would also like to thank the National BioResource Project (Japan) for the *lgc-54* mutant strains (FX3448 and FX3518).

## Supporting information captions

**S1 Fig. Dose response with four divergent wild isolates.** Results from the abamectin dose response HTA for brood size (norm.n), animal length (mean.TOF), animal optical density (mean.EXT), and normalized optical density (mean.norm.EXT) are shown. For each trait, drug concentration (nM) (x-axis) is plotted against phenotype subtracted from control (y-axis) and colored by strain (CB4856: blue, DL238: green, JU775: purple, N2: orange). A concentration of 7.5 nM was chosen for future experiments.

**S2 Fig. Genome-wide association mappings identify six QTL across three traits in response to abamectin. A)** Normalized residual phenotype (y-axis) of 210 wild isolates (x-axis) in response to abamectin. **B)** Association mapping results are shown. Genomic position (x-axis) is plotted against the −log10*(p)* value (y-axis) for each SNV. SNVs are colored pink if they pass the genome-wide eigen-decomposition significance threshold designated by the grey line. The genomic regions of interest that pass the significance threshold are highlighted by pink rectangles. **C)** For each QTL, the normalized residual phenotype (y-axis) of strains split by genotype at the peak marker (x-axis) are plotted as Tukey box plots. Each point corresponds to a wild isolate strain. Strains with the N2 reference allele are colored grey, and strains with an alternative allele are colored pink.

**S3 Fig. Linkage mapping identifies 14 QTL across four traits in response to abamectin. A)** Normalized residual phenotype (y-axis) of 225 RIAILs (x-axis) in response to abamectin. The parental strains are colored: N2, orange; CB4856, blue. **B)** Linkage mapping results are shown. Genomic position in Mb (x-axis) is plotted against the logarithm of the odds (LOD) score (y-axis) for 13,003 genomic markers. Each significant QTL is indicated by a red triangle at the peak marker, and a blue rectangle shows the 95% confidence interval around the peak marker. The percentage of the total variance in the RIAIL population that can be explained by each QTL is shown above the QTL. **C)** For each QTL, the normalized residual phenotype (y-axis) of RIAILs split by genotype at the marker with the maximum LOD score (x-axis) are plotted as Tukey box plots. Each point corresponds to a unique recombinant strain. Strains with the N2 allele are colored orange, and strains with the CB4856 allele are colored blue.

**S4 Fig. Summary of QTL mapping for responses to abamectin.** Genomic positions (x-axis) of all QTL identified from linkage mapping (top) and association mapping (bottom) are shown for each drug-trait (y-axis). Each QTL is plotted as a point at the genomic location of the peak marker and a line that represents the confidence interval. QTL are colored by the significance of the LOD score (linkage) or −log10(*p*) value (association), increasing from purple to green to yellow.

**S5 Fig. Two-dimensional genome scan for mean optical density (mean.EXT) in abamectin.** Log of the odds (LOD) scores are shown for each pairwise combination of loci, split by chromosome. The upper-left triangle contains the epistasis LOD scores (interaction effects), and the lower-right triangle contains the LOD scores for the full model (both interaction and additive effects). LOD scores are colored by significance, increasing from purple to green to yellow. The LOD scores for the epistasis model are shown on the left of the color scale, and the LOD scores for the full model are shown on the right.

**S6 Fig. Chromosome substitution strains validate the existence of one or more resistance loci on chromosome V. A)** Strain genotypes are shown as colored rectangles (N2: orange, CB4856: blue) in detail for chromosome V (left) and in general for the rest of the chromosomes (right). **B)** Normalized residual mean lengths in abamectin (mean.TOF, x-axis) are plotted as Tukey box plots against strain (y-axis). Statistical significance of each NIL as compared to its parental strain (ECA573 to N2 and ECA554 to CB4856) calculated by Tukey’s HSD is shown above each strain (ns = non-significant (p-value > 0.05); *, **, ***, and *** = significant (p-value < 0.05, 0.01, 0.001, or 0.0001, respectively).

**S7 Fig. Refining QTL positions with NILs. A)** Fine mapping of all common variants on chromosome V is shown. Genomic position (x-axis) is plotted against the −log10(*p*) values (y-axis) for each variant and colored by the genotype of the variant in the CB4856 strain (grey = N2 reference allele, blue = variation from the N2 reference allele). Genomic regions identified from linkage mapping analysis are highlighted in blue and genomic regions identified from association mapping are highlighted in pink. The horizontal grey line represents the genome-wide eigen-decomposition significance threshold. The red points represent the positions of the most significant variants in the genes *glc-1* (diamond), *glc-3* (circle), and *lgc-54* (square). The vertical lines represent the smallest NIL-defined genomic region for the VL (solid), VC (dashed), and VR (dotted) QTL. **B)** Strain genotypes are shown as colored rectangles (N2: orange, CB4856: blue) in detail for chromosome V. The vertical lines represent the smallest NIL-defined genomic region for the VL (solid), VC (dashed), and VR (dotted) QTL.

**S8 Fig. Mediation analysis for the VL and VC QTL.** Mediation estimates calculated as the indirect effect that differences in expression of each gene plays in the overall phenotype (y-axis) are plotted against genomic position of the eQTL (x-axis) on chromosome V for all genes with a gene expression QTL in the narrowed VL and VC intervals. The 90th percentile of the distribution of mediation estimates is represented by the horizontal grey line. The confidence of the estimate increases (*p*-value decreases) as points become more solid.

**S9 Fig. NILs validate and narrow the VC QTL. A)** Strain genotypes are shown as colored rectangles (N2: orange, CB4856: blue) in detail for chromosome V (left) and in general for the rest of the chromosomes (right). The dashed vertical lines represent the previous NIL-defined QTL interval for VC. **B)** Normalized residual mean optical densities in abamectin (mean.EXT, x-axis) are plotted as Tukey box plots against strain (y-axis). Statistical significance of each NIL as compared to the N2 strain calculated by Tukey’s HSD is shown above each strain (ns = non-significant (p-value > 0.05); *, **, ***, and *** = significant (p-value < 0.05, 0.01, 0.001, or 0.0001, respectively).

**S1 File. Dose response phenotype data.** Results of the dose response for the genome-wide association high-throughput fitness assay

**S2 File. Wild isolate phenotype data.** Residual phenotypic values for the 210 wild isolates in response to abamectin

**S3 File. Association mapping results.** Genome-wide association mapping results for all four drug-response traits tested in the high-throughput fitness assay

**S4 File. RIAIL phenotype data.** Residual phenotypic values for the 225 set 2 RIAILs in response to abamectin

**S5 File. Linkage mapping results.** Linkage mapping LOD scores at 13,003 genomic markers for all four drug-response traits with the set 2 RIAILs

**S6 File. Summary of two-dimensional genome scan.** Summary of the scan2 object containing data from the two-dimensional genome scan with animal optical density (mean.EXT) in abamectin

**S7 File. Chromosome V variants.** Correlation values and annotations for all variants on chromosome V

**S8 File. NIL sequence data.** VCF from the whole-genome sequencing for all the NILs in this study

**S9 File. NIL genotype data.** Simplified genotypes of the NILs in the study

**S10 File. CSSV phenotype data.** Raw pruned phenotypes for the high-throughput fitness assay with the chromosome V substitution strains

**S11 File. Statistical significance for NIL assays.** Pairwise statistical significance for all strains and high-throughput assays

**S12 File. ChrV NIL breakup phenotype data.** Raw pruned phenotypes for the NILs used to break up the QTL interval on chromosome V

**S13 File. Candidate genes on chrV.** List of all genes in the chromosome VL and VC intervals, their functional descriptions and GO annotations, and if they have variation in CB4856

**S14 File. Set 1 RIAIL phenotype data.** Residual phenotypic values for the 107 set 1 RIAILs in response to abamectin

**S15 File. Mediation estimates for chrV QTL.** Mediation estimates for the chromosome VL and VC QTL

**S16 File. NILs to narrow VC QTL.** Raw pruned phenotypes for the chromosome VC NIL high-throughput fitness assay

**S17 File. lgc-54 mutant phenotypes.** Raw pruned phenotypes for the the lgc-54 deletion strains high-throughput fitness assay

**S18 File. One-to-one orthologous genes in** *H. contortus*. Coordinates and orthologous relationships for *H. contortus* QTL

**S19 File. Orthogroups for** *H. contortus* **and** *C. elegans* **genes.** Contains a list of all the genes found in each orthogroup between *C. elegans* and *H. contortus*

